# Development of a high-throughput, quantitative platform using human cerebral organoids to study virus-induced neuroinflammation in Alzheimer’s disease

**DOI:** 10.1101/2024.03.21.585957

**Authors:** Meagan N. Olson, Pepper Dawes, Liam F. Murray, Nathaniel J. Barton, Jonathan Sundstrom, Adrian R. Orszulak, Samantha M. Chigas, Khanh Tran, Aimee J. Aylward, Michele F. Caliandro, Sean-Patrick H. Riechers, Khashayar Afshari, Qi Wang, Manuel Garber, Fiachra Humphries, Megan H. Orzalli, Douglas T. Golenbock, Michael T. Heneka, Hyung Suk Oh, George M. Church, Tracy L. Young-Pearse, David M. Knipe, Benjamin Readhead, Yingleong Chan, Elaine T. Lim

## Abstract

Neuroinflammation is a central process in the pathogenesis of several neurodegenerative diseases such as Alzheimer’s disease (AD), and there are active efforts to target pathways involved in neuroinflammation for molecular biomarker discovery and therapeutic development in neurodegenerative diseases. It was also proposed that there may be an infectious etiology in AD that is associated with viruses such as herpes simplex virus (HSV-1) and influenza A virus (IAV), leading to neuroinflammation-induced AD pathogenesis or disease progression. We sought to develop high-throughput, quantitative molecular biomarker assays using dissociated cells from human cerebral organoids (dcOrgs), that can used for screening compounds to reverse AD-associated neuroinflammation. We found that HSV-1 infection, but not IAV infection, in dcOrgs led to increased intracellular Aβ42 and phosphorylated Tau-Thr212 (pTau-212) expression, lower ratios of secreted Aβ42/40, as well as neuronal loss, and increased proportions of astrocytes and microglia, which are hallmarks of AD. Among the glia cell-type markers, Iba1 (microglia) and GFAP (astrocyte) expression were most strongly correlated with HSV-1 expression, which further supported that these biomarkers are perturbed by glia-mediated neuroinflammation. By performing large-scale RNA sequencing, we observed that differentially expressed transcripts in HSV-1 infected dcOrgs were specifically enriched for AD-associated GWAS genes, but not for genes associated with other common neurodegenerative, neuropsychiatric or autoimmune diseases. Immediate treatment of HSV-1 infected dcOrgs with anti-herpetic drug acyclovir (ACV) rescued most of the cellular and transcriptomic biomarkers in a dosage-dependent manner, indicating that it is possible to use our high-throughput platform to identify compounds or target genes that can reverse these neuroinflammation-induced biomarkers associated with AD.

## INTRODUCTION

There is mounting evidence that chronic neuroinflammation mediated by microglia and astrocytes can lead to Alzheimer’s disease (AD) pathogenesis, disease progression and severity^1,2^. This evidence has led to active research on nonsteroidal anti-inflammatory drugs and modulation of key neuroinflammation targets for AD therapeutics, such as NLRP3 inflammasome, TREM2 and INPP5D^3–7^. It was also proposed that there is an infectious etiology in AD and viruses such as the double-stranded DNA herpes simplex virus (HSV-1) and single-stranded RNA influenza A virus (IAV) can trigger neuroinflammation that may lead to AD pathogenesis, progression or severity^8–13^.

Epidemiological studies on large-scale population cohorts further supported the infectious hypothesis in AD through the discoveries that viral encephalitis and influenza were associated with increased risk for AD and anti-herpetic medication reduced dementia risk in patients with HSV-1 infections^14–20^. The anti-herpetic drug acyclovir (ACV) was shown to be highly effective in reducing HSV-1 replication and reducing clinical symptoms^21^. Clinical trials using the oral anti-herpetic drug valacyclovir (VCV) as a potential treatment for AD-associated neuroinflammation are currently underway and VCV may prove to be a promising AD therapeutic^22–25^.

In parallel, several animal and human *in-vitro* models had been developed for AD research, including human induced pluripotent stem cell (hiPSC) derived models such as two dimensional (2D) induced neurons and three dimensional (3D) brain organoids^26–30^, to advance biomarker and therapeutics discovery for AD. The brain organoid models contain significant proportions of cell types that are more commonly found in human fetal brains than adult brains, such as neural progenitor cells^31–34^, and therefore, are imperfect models for neurodegenerative diseases. However, despite being imperfect, the use of human brain organoids had already led to novel, exciting molecular insights into AD neuropathology^28,35^. In recent years, researchers discovered that HSV-1 catalyzed β-amyloid (Aβ) aggregation and led to encephalitis-associated cell-type specific aberrations in human brain organoids, thus illuminating molecular mechanisms that might be involved in the infectious etiology of AD^36–44^.

Given these exciting prior discoveries, we sought to identify molecular and cellular biomarkers to develop high-throughput, quantitative assays around the use of human cerebral organoids, specifically for virus-induced neuroinflammation in AD. Microglia, the resident immune cells in the central nervous system, are differentiated from the mesodermal lineage *in-vitro*^45,46^. As such, it was widely thought that cerebral organoids did not contain microglia and would not produce interferon response upon viral infection. However, earlier studies had discovered and characterized CD11b+ microglia that differentiated innately within these cerebral organoids^47–50^.

Building on top of these extensive prior research and technological development, we aim to develop a high-throughput, quantitative human *in-vitro* platform that can be used to identify novel targets and compounds to reverse neuroinflammation-induced molecular and cellular AD-associated neuropathology.

## RESULTS

### 3D cerebral organoids versus 2D dissociated cells from cerebral organoids

We differentiated several batches of 3D cerebral organoids from a control donor without AD using a previously established protocol^30,34^. We first performed HSV-1 (that was tagged with green fluorescent protein GFP) viral infections in intact 3D cerebral organoids that were differentiated for 3 months, and found that only ∼5% of cells were infected and the infected cells were primarily on the periphery of the organoids (Figure S1A). The low fractions of infected cells, as well as non-uniform infection of the 3D cerebral organoids, would pose challenges for a neuroinflammation model in AD.

As such, we dissociated cells from 2-month to 4-month 3D cerebral organoids for replating as 2D monolayer cell cultures, followed by recovery of the dissociated cells for a month. We referred to dissociated cells from the cerebral organoids as dcOrgs in this paper. We performed HSV-1 infections in dcOrgs and consistently observed >50% of cells were infected.

In addition, given that microglia is an essential immune cell type in a neuroinflammation model, we evaluated the fractions of dcOrgs that are positive for P2RY12 expression (homeostatic microglia), Iba1 expression (activated microglia) and GLAST expression (astrocytes). We found that there were low fractions of Iba1+ microglia (0.02%) in dcOrgs, and 7.1% of dcOrgs were P2RY12+ microglia (Figure S1B), consistent with reported fractions found in human and mouse brains^51^. We also found that 3.8% of dcOrgs were GLAST+ astrocytes, which was lower than reported fractions found in human adult brains^52–54^.

There are several advantages in using dcOrgs as a model for neuroinflammation-induced AD neuropathology, instead of 3D cerebral organoids, such as the ability to obtain higher percentages of infected cells after a day post-inoculation, more uniform distribution of viral infections, and dcOrgs can be frozen down for longer-term storage and passaged as regular 2D cell cultures to obtain large quantities of cells for high-throughput screens. Both HSV-1 and IAV infections had been associated with increased risk for AD^19^, although HSV-1 is a neurotropic DNA virus and IAV is a respiratory RNA virus. Subsequently, we used dcOrgs for HSV-1 and IAV infections to identify if there may be potential neuroinflammation-induced molecular or cellular biomarkers associated with AD.

### HSV-1 infection and ACV treatment induced major human transcriptome perturbations in dcOrgs but not in hiPSCs

Initially, we conducted 3 sets of experiments to compare HSV-1 infected dcOrgs versus uninfected dcOrgs (*Inf-vs-Ctrl dcOrgs*), HSV-1 infected dcOrgs versus HSV-1 infected and ACV-treated dcOrgs (*Inf-vs-ACV dcOrgs*), and HSV-1 infected dcOrgs versus dcOrgs with UV-inactivated HSV-1 added (*Inf-vs-UV dcOrgs*), shown in Figures 1A-B. The first 2 sets of experiments (*Inf-vs-Ctrl dcOrgs* and *Inf-vs-ACV dcOrgs*) were repeated for replication. In addition, we conducted another 2 sets of experiments to compare HSV-1 infected hiPSCs from the same donor versus uninfected hiPSCs (*Inf-vs-Ctrl hiPSCs*) and HSV-1 infected hiPSCs versus HSV-1 infected and ACV-treated hiPSCs (*Inf-vs-ACV hiPSCs*). Each set of experiments was performed in triplicates, followed by total RNA extraction, ribosomal RNA depletion and RNA sequencing (RNA-seq).

**Figure 1:**
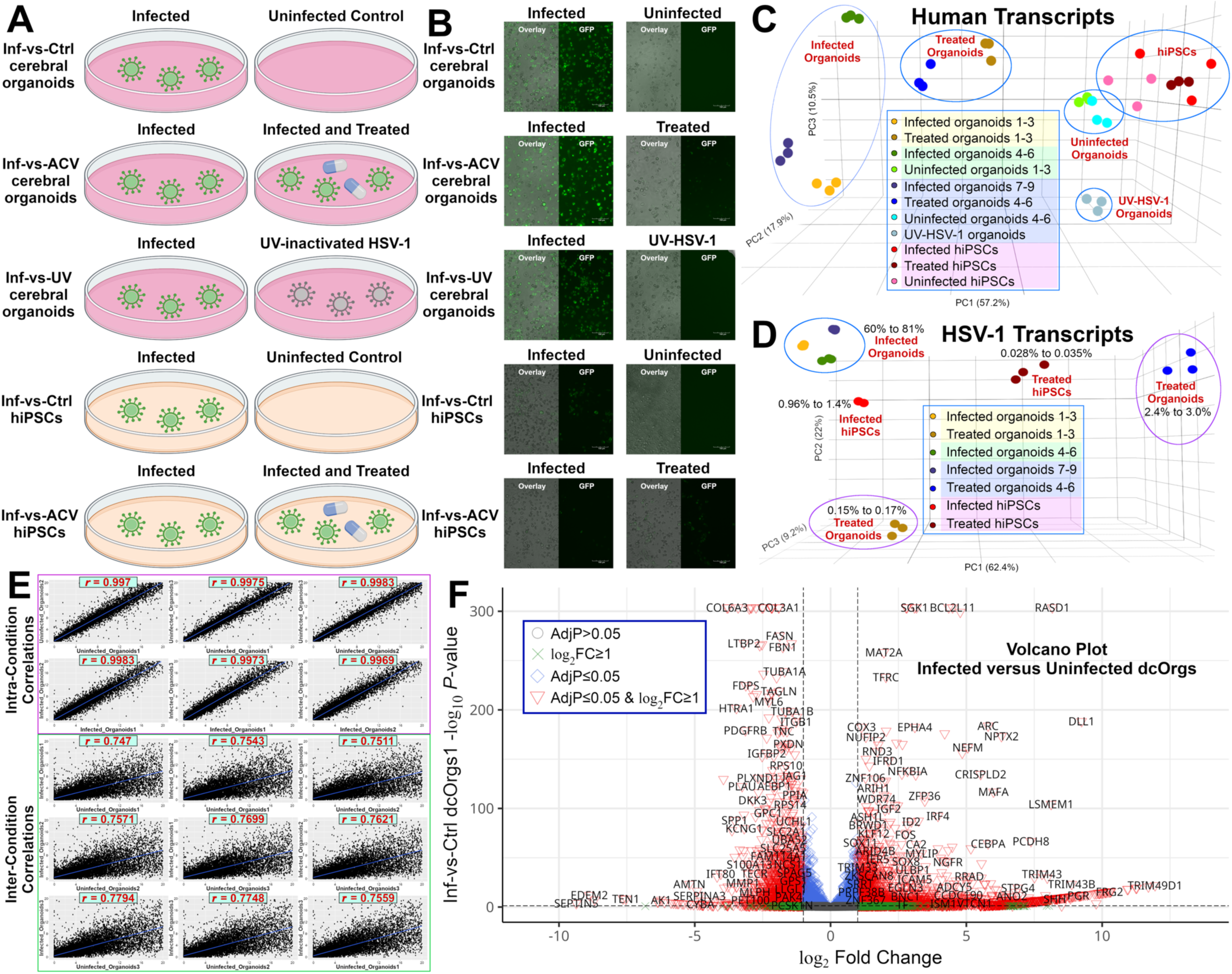
Experimental setup and overview of the results from the RNA sequence datasets. (A) Five sets of experimental conditions that our study generated RNA-seq data on (*Inf-vs-Ctrl dcOrgs*, *Inf-vs-ACV dcOrgs*, *Inf-vs-UV dcOrgs*, *Inf-vs-Ctrl hiPSCs*, *Inf-vs-ACV hiPSCs*). (B) Fluorescence images (20X magnification) of the cells across the experimental conditions in our study. (C) 3D PCA plot of the human transcripts (in FPKM), with the colored tints in the legend representing the experimental batches. Each number (1-9) represents an independent biological sample. (D) 3D PCA plot of the HSV-1 transcripts (in FPKM), with the colored tints in the legend representing the experimental batches. Each number (1-9) represents an independent biological sample. The percentages indicate the percentages of total transcripts that mapped to the HSV-1 transcriptome for each group of samples. The two sets of ACV-treated organoids are circled in purple and the infected organoids are circled in blue. (E) Intra-condition and inter-condition correlation plots showing pairwise FPKM values and Pearson’s *r* for *Inf-vs-Ctrl dcOrgs1* (Infected_Organoids4-6 versus Uninfected_Organoids1-3). (F) Volcano plot of differentially expressed human genes for *Inf-vs-Ctrl dcOrgs1* (Infected_Organoids4-6 versus Uninfected_Organoids1-3), with the *log_2_* fold change calculated as the expression of the transcript from HSV-1 infected dcOrgs versus uninfected dcOrgs, and - *log_10_ P*-values were calculated using the Benjamini-Hochberg adjusted *P*-values of the differential expression.

3D principal components analysis (PCA) showed that most of the variation (57.2%) in the human transcripts was captured by the first principal component (PC1) (Figure 1C, Figure S2A). The dcOrgs were also distinctively separated by their conditions into well-defined clusters on PC1. dcOrgs with UV-inactivated HSV-1 were clustered close to uninfected dcOrgs on PC1, whereas ACV-treated dcOrgs formed their own cluster between infected and uninfected dcOrgs on PC1. All hiPSCs clustered together into a single group regardless of the condition (infected, uninfected or ACV-treated).

To elucidate the transcriptome-wide expression patterns of HSV-1, we similarly performed PCA using detected HSV-1 transcripts and found that most of the variation (62.4%) was captured by PC1 (Figures 1D, Figure S2B). All of the infected dcOrgs clustered together and infected hiPSCs clustered closely with the infected dcOrgs. The ACV-treated hiPSCs and dcOrgs formed their own clusters, separate from the infected samples. However, both sets of ACV-treated dcOrgs showed distinctive clusters, primarily in PC1 and PC2. We found that 0.15-0.17% of all transcripts were viral transcripts in the first set of ACV-treated dcOrgs (dcOrgs1), 2.4-3% of total transcripts were viral transcripts in the second set of ACV-treated dcOrgs (dcOrgs2), whereas 60-81% of all transcripts were viral transcripts in infected dcOrgs. There were 14-20 times more HSV-1 transcripts in the second set of ACV-treated dcOrgs compared to the first set of ACV-treated dcOrgs, suggesting that ACV treatment in the second set of dcOrgs was not as effective as in the first set of dcOrgs on suppressing viral transcript expression. As such, both sets of ACV-treated dcOrgs can inform us about viral dosage-dependent effects on human transcript expression.

A fraction (0.96-1.4%) of all transcripts were viral transcripts in infected hiPSCs, and 0.028-0.035% of all transcripts were viral transcripts in ACV-treated hiPSCs. There were lower viral transcript counts in the hiPSCs than the dcOrgs, despite using a higher multiplicity of infection (MOI) for the hiPSCs versus the dcOrgs (MOI of 4 and 2 respectively), consistent with prior reports that stem cells can be highly resistant to viral infection, unlike differentiated cells^55,56^.

### High inter-sample correlations observed in dcOrgs and hiPSCs

Similar to our previous study^34^, we observed high correlations between biological replicates of dcOrgs by pooling large numbers of 3D cerebral organoids for the initial dissociation. Pairs of infected dcOrg replicates or pairs of uninfected dcOrg replicates had high correlations and low variability (Pearson’s *r*>0.99; Figure 1E, Table S1). The correlations between infected-uninfected dcOrg pairs were lower (*r*=0.75-0.82). The correlations between infected-treated dcOrg pairs ranged from *r*=0.81-0.89. The correlations between infected-uninfected or infected-treated hiPSC pairs were higher (*r*=0.95-1), possibly because fewer human transcripts were differentially expressed from HSV-1 infection in hiPSCs compared to dcOrgs.

### HSV-1 infection preferentially up-regulated human transcripts in dcOrgs and hiPSCs that was not restored by ACV treatment

The DEG analyses of human transcripts from HSV-1 infected versus uninfected dcOrgs showed that there were more up-regulated human transcripts compared to down-regulated human transcripts (odds ratios, *OR*=1.13 and 1.14; family-wise error rate, *FWER*=5.9×10^-4^ and 1.2×10^-4^; Figure 1F, Figure S3A, Tables S2-3). Similarly, there were more up-regulated human transcripts in HSV-1 infected dcOrgs versus dcOrgs with UV-inactivated HSV-1 (*OR*=1.21, *FWER*=5.8×10^-9^; Figure S3B). In hiPSCs, we observed a similar imbalance with more up-regulated human transcripts in HSV-1 infected versus uninfected hiPSCs (*OR*=1.3, *FWER*=5.0×10^-4^; Figure S3C).

There were equal proportions of up-regulated or down-regulated human transcripts in HSV-1 infected versus ACV-treated dcOrgs (*OR*=1.02 and 1.03, *FWER*=0.42 and 0.36 respectively; Figures S3D-E) and in ACV-treated versus uninfected dcOrgs and ACV-treated versus dcOrgs with UV-inactivated HSV-1 (*OR*=1.1 and 1.15, *FWER*=0.16 and 0.051 respectively; Figures S3F-H). However, there was a significant excess of up-regulated human transcripts observed in HSV-1 infected versus ACV-treated hiPSCs (*OR*=9.9, *FWER*=3.4×10^-7^; Figure S4A). No imbalances in the proportions of up-regulated or down-regulated viral transcripts in HSV-1 infected versus ACV-treated dcOrgs or hiPSCs were observed (Figures S4B-D, Tables S3-4).

Collectively, these results showed that HSV-1 infection preferentially up-regulated the expression of human transcripts in dcOrgs, indicating that there are proportionally more human transcripts and pathways that can be attenuated to reduce the expression of these up-regulated transcripts in HSV-1 infected dcOrgs, versus to increase the expression of down-regulated transcripts in HSV-1 infected dcOrgs. However, ACV treatment on infected dcOrgs did not significantly rescue these global imbalances (Figure S3H), indicating that the use of anti-viral drug ACV is not sufficient for a significant reversal of HSV-1 induced transcriptomic changes in dcOrgs.

### HSV-1 expression, but not IAV expression, in dcOrgs was highly positively correlated with intracellular expression of Aβ42 and phosphorylated Tau-Thr212 (pTau-212)

Given that HSV-1 infection preferentially up-regulated the expression of many human transcripts in dcOrgs, we wondered if these transcriptomic perturbations may affect the expression of intracellular Aβ1-42 (Aβ42) or phosphorylated Tau on residue Thr212 (pTau-212). Several prior studies have shown that accumulation of intracellular Aβ (intracellular Aβ42 in particular) precedes the accumulation of extracellular Aβ plaques^57^. We performed 3-channel flow cytometry with 20,000-30,000 single cells for each condition, followed by machine learning based gating (Figure 2A). A Zombie Violet dye was used to label dead cells, Alexa Fluor 647 or Allophycocyanin (APC) fluorophore-conjugated antibodies were used to detect the expression of Aβ42, pTau-212 or cell-type specific markers, and the HSV-1 virus was tagged with GFP. One advantage of flow cytometry is the ability to quantitatively measure the intensities of intracellular Aβ42/pTau-212 and HSV-1 for each single cell. This enables us to directly evaluate if there may be higher expression of intracellular Aβ42/pTau-212 in infected versus uninfected cells.

**Figure 2:**
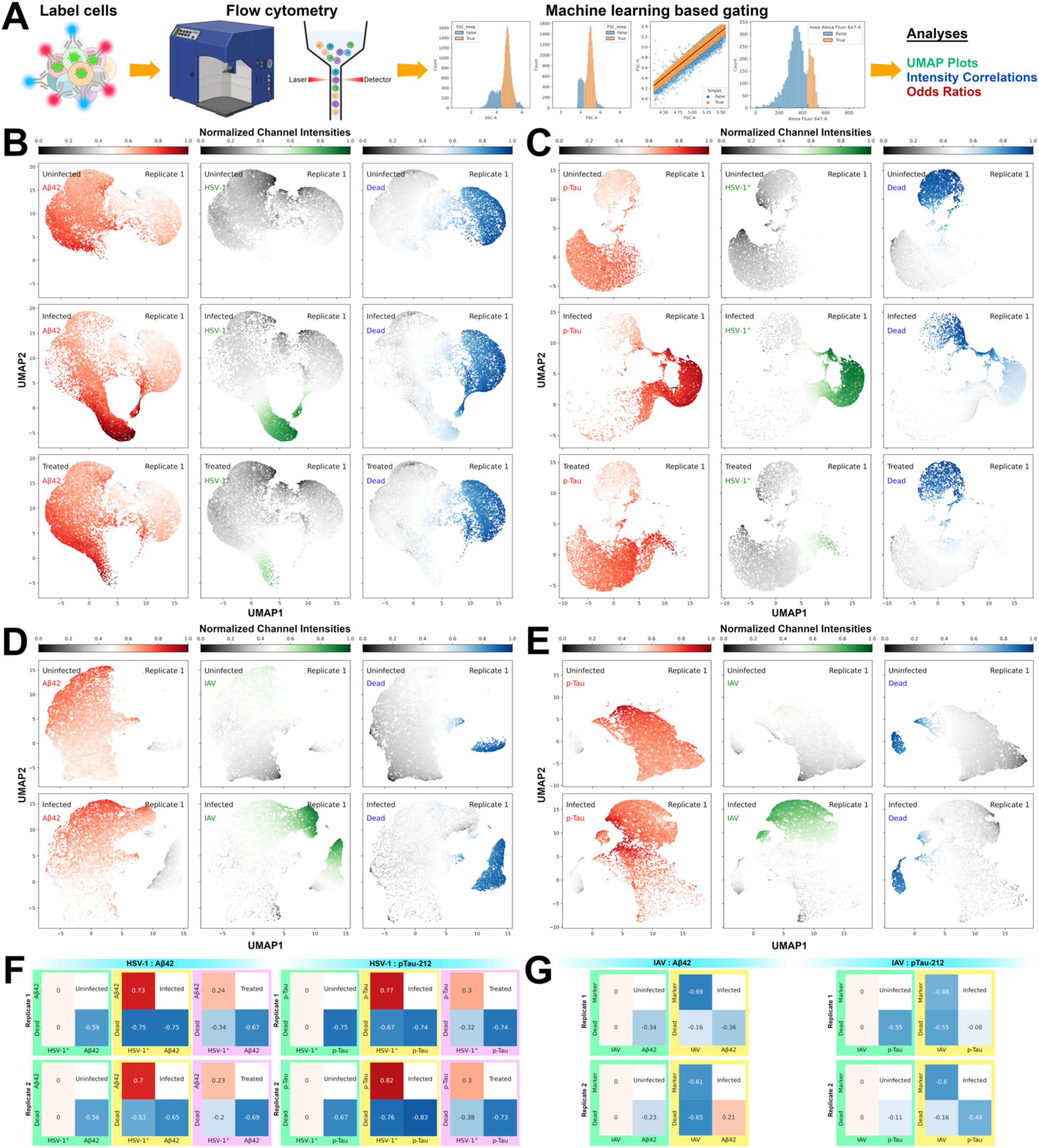
Co-expression of HSV-1 and IAV with intracellular Aβ42 and pTau-212. (A) Schematic of the flow cytometry experiments and computational analyses. (B) UMAP plots showing the normalized fluorophore intensities for Aβ42 (red), HSV-1 (green) and Zombie fluorophore (blue) across uninfected, HSV-1 infected and ACV-treated cells. (C) UMAP plots showing the normalized fluorophore intensities for pTau-212 (red), HSV-1 (green) and Zombie fluorophore (blue) across uninfected, HSV-1 infected and ACV-treated cells. (D) Pairwise adjusted correlation heatmaps using the intensities for HSV-1, Zombie fluorophore and Aβ42 or pTau-212 across uninfected, HSV-1 infected and ACV-treated cells. The adjusted correlations for 2 sets of replicates (Replicate 1 and Replicate 2) were shown. (E) UMAP plots showing the normalized fluorophore intensities for Aβ42 (red), IAV (green) and Zombie fluorophore (blue) across uninfected and IAV infected cells. (F) UMAP plots showing the normalized fluorophore intensities for pTau-212 (red), IAV (green) and Zombie fluorophore (blue) across uninfected and IAV infected cells. (G) Pairwise adjusted correlation heatmaps using the intensities for IAV, Zombie fluorophore and Aβ42 or pTau-212 across uninfected and IAV-infected cells. The adjusted correlations for 2 sets of replicates (Replicate 1 and Replicate 2) were shown.

Uniform Manifold Approximation and Projection (UMAP) plots revealed that there were higher intracellular Aβ42 and pTau-212 expression in HSV-1 infected dcOrgs compared to uninfected dcOrgs or ACV-treated dcOrgs (Figures 2B-C). We next asked if IAV infection in dcOrgs could similarly lead to increased intracellular Aβ42 and pTau-212 expression. In contrast, we found that IAV infection of dcOrgs did not result in higher expression of Aβ42 and pTau-212 (Figures 2D-E).

Next, we directly evaluated if there were higher Aβ42 or pTau-212 expression in cells with higher expression (or more copies) of HSV-1 or IAV. We calculated the adjusted correlations in intensities and observed a high correlation in Aβ42 and HSV-1 intensities (*r*=0.73 and 0.7 for 2 replicates; Figure 2F), and ACV treatment reduced the correlation in Aβ42 and HSV-1 intensities (*r*=0.24 and 0.23; Figure 2F). Correlations in the intensities of pTau-212 and HSV-1 were similarly high (*r*=0.77 and 0.82; Figure 2F) and ACV treatment similarly lowered the correlation in intensities of pTau-212 and HSV-1 (*r*=0.3 for both replicates; Figure 2F). In contrast, we observed negative correlations in Aβ42 and IAV intensities (*r*=-0.61 and -0.65; Figure 2G), as well as pTau-212 and IAV intensities (*r*=-0.48 and -0.6; Figure 2G). The amyloid precursor protein (APP) that results in Aβ peptides after proteolytic cleavage, is primarily expressed in neurons^58^. As such, the negative correlations in Aβ42/pTau-212 and IAV intensities may indicate that the cells that facilitated IAV replication in dcOrgs were primarily non-neuronal cells.

These results showed that HSV-1 infection in dcOrgs led to increased intracellular expression of Aβ42 and pTau-212, but IAV infection in dcOrgs did not. In addition, intracellular expression of Aβ42 and pTau-212 were positively strongly correlated with intracellular expression of HSV-1, but not with intracellular expression of IAV.

### High correlations in intracellular Aβ42 and HSV-1 expression were not predominantly driven by Aβ monomers

We evaluated if the high correlations in intracellular Aβ42 and HSV-1 expression that we had observed may be driven by intracellular Aβ monomers, by using the solanezumab monoclonal antibody (Figure 3A). Solanezumab was recently reported to have failed to slow cognitive decline in people with preclinical AD or mild dementia due to AD^59,60^. We found positive but weaker correlations in the expression of intracellular Aβ monomers and HSV-1 (*r*=0.38 and 0.46; Figure 3B) and ACV treatment on HSV-1 infected dcOrgs reduced the correlations (*r*=0.18 and 0.25; Figure 3B). These results indicate that while the expression of HSV-1 in dcOrgs is correlated with the expression of Aβ monomers, the stronger correlations that we had observed in co-expression of intracellular Aβ42 and HSV-1 were not likely to be predominantly driven by Aβ monomers.

**Figure 3:**
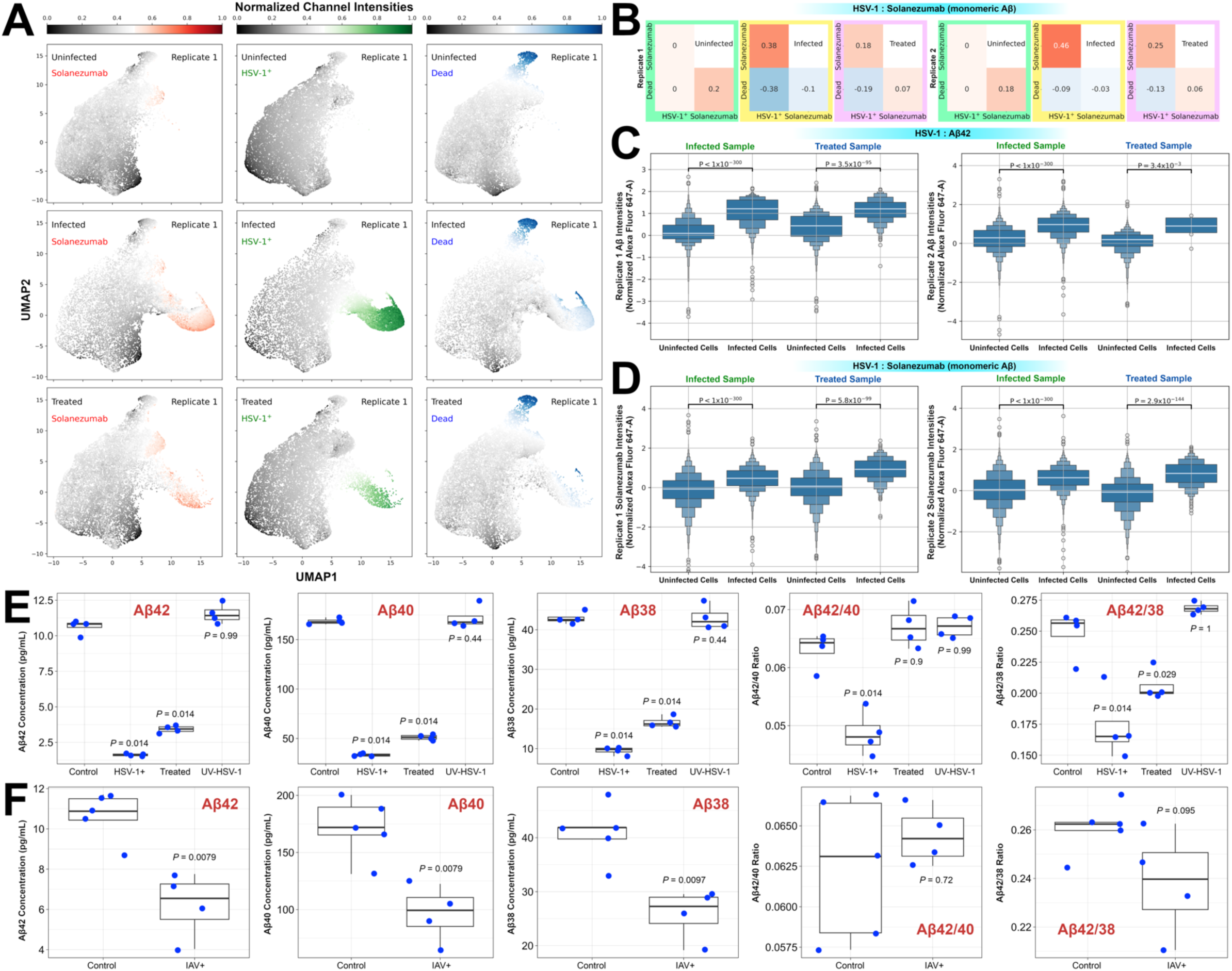
Evaluating Aβ monomers and extracellular Aβ42/40/38 concentrations. (A) UMAP plots showing the normalized fluorophore intensities for Aβ monomers using Solanezumab (red), HSV-1 (green) and Zombie fluorophore (blue) across uninfected, HSV-1 infected and ACV-treated cells. (B) Pairwise adjusted correlation heatmaps using the intensities for HSV-1, Zombie fluorophore and Solanezumab (Aβ monomers) across uninfected, HSV-1 infected and ACV-treated cells, shown for 2 sets of replicates (Replicate 1 and Replicate 2). (C) Boxen plots showing the normalized Aβ42 fluorophore intensities for uninfected cells (HSV-1^-^ cells) versus infected cells (HSV-1^+^ cells) within the same infected sample, as well as uninfected cells versus infected cells within the same ACV-treated sample. (D) Boxen plots showing the normalized Solanezumab fluorophore intensities for uninfected cells (HSV-1^-^ cells) versus infected cells (HSV-1^+^ cells) within the same infected sample, as well as uninfected cells versus infected cells within the same ACV-treated sample. (E) Concentrations of Aβ42/40/38 in pg/mL, Aβ42/40 ratios and Aβ42/38 ratios detected from conditioned media that had undergone heat inactivation for uninfected (control) dcOrgs, HSV-1 infected (HSV-1+) dcOrgs, HSV-1 infected and ACV-treated (Treated) dcOrgs, and dcOrgs with UV-inactivated HSV-1 (UV-HSV-1). *P*-values shown were calculated using 1-sided Wilcoxon ranked sum test with comparison to control uninfected dcOrgs. (F) Concentrations of Aβ42/40/38 in pg/mL, β42/40 ratios and Aβ42/38 ratios detected from conditioned media that had undergone heat inactivation for uninfected (control) dcOrgs and IAV infected (IAV+) dcOrgs. *P*-values shown were calculated using 1-sided Wilcoxon ranked sum test with comparison to control uninfected dcOrgs.

### Direct comparisons in uninfected versus infected cells within the same samples validate higher intracellular expression of Aβ42 and Aβ monomers

Since not all cells within our HSV-1 infected dcOrgs or ACV-treated dcOrgs were infected, we evaluated the intensity distributions of intracellular Aβ42 from uninfected cells versus infected cells within the same HSV-1 infected samples (Figure 3C) and found that the intracellular Aβ42 intensity distributions from infected cells were significantly higher than the distributions from uninfected cells (1-tailed Wilcoxon ranked sum *P*<1×10^-300^ for both replicates). The intensity distributions from infected cells were also significantly higher than the distributions from uninfected cells within the same ACV-treated samples (1-tailed Wilcoxon tanked sum *P*<3.5×10^-95^ and *P*<3.4×10^-3^ for both replicates).

Similarly, the solanezumab intensity distributions from infected cells were significantly higher than the distributions from uninfected cells within the same HSV-1 infected samples (Figure 3D; 1-tailed Wilcoxon ranked sum *P*<1×10^-300^ for both replicates), as well as within the same ACV-treated samples (1-tailed Wilcoxon tanked sum *P*<5.8×10^-99^ and *P*<2.9×10^-144^ for both replicates). These results provide direct validation for our correlation analyses and support the observations that HSV-1 led to increased intracellular expression of Aβ42 and Aβ monomers.

### HSV-1 infection significantly lowered extracellular Aβ42/40 ratios detected in conditioned media but IAV infection did not affect extracellular Aβ42/40 ratios

To follow up on our observations that HSV-1 infection on dcOrgs led to increased accumulation of intracellular Aβ42 and pTau-212 that could be rescued by immediate ACV treatment, we tested if HSV-1 infection in dcOrgs may affect the concentrations of secreted Aβ42, Aβ40 or Aβ38 peptides detected in conditioned media, and if immediate ACV treatment may restore the concentrations of secreted Aβ42, Aβ40 or Aβ38 peptides. We performed high-throughput ELISA assays to evaluate the concentrations of extracellular Aβ peptides in conditioned media collected from dcOrgs that were uninfected, infected with HSV-1, infected and treated with ACV, as well as dcOrgs with UV-inactivated HSV-1 added. We performed heat inactivation on the conditioned media prior to conducting the ELISA assays.

Conditioned media from HSV-1 infected and ACV-treated dcOrgs had consistently lower concentrations of extracellular Aβ42, Aβ40 and Aβ38 compared to control conditioned media from uninfected dcOrgs (1-tailed Wilcoxon *P*=0.014; Figure 3D, Table S5). There were significantly lower extracellular Aβ42/40 and Aβ42/38 ratios in conditioned media from HSV-1 infected dcOrgs (1-tailed Wilcoxon *P*=0.014; Figure 3D). ACV treatment restored extracellular Aβ42/40 ratios to similar ratios as detected in control conditioned media from uninfected dcOrgs (1-tailed Wilcoxon *P*=0.9) but did not completely restore extracellular Aβ42/38 ratios (1-tailed Wilcoxon *P*=0.029).

IAV infection in dcOrgs similarly led to decreased concentrations of extracellular Aβ42, Aβ40 and Aβ38 detected in the conditioned media, compared to control conditioned media from uninfected dcOrgs (1-tailed Wilcoxon *P*=0.0097; Figure 3E). However, extracellular Aβ42/40 and Aβ42/38 ratios in conditioned media from IAV infected dcOrgs did not differ significantly from ratios detected in control conditioned media from uninfected dcOrgs (1-tailed Wilcoxon *P*=0.72 and 0.095 respectively; Figure 3E). We repeated the ELISA assays using conditioned media that were subjected to UV-radiation and obtained similar results (Table S5, Figure S5).

These results showed that HSV-1 infection in dcOrgs resulted in significantly lower ratios of extracellular Aβ42/40, but IAV infection in dcOrgs did not affect the ratios of extracellular Aβ42/40. In addition, ACV treatment of HSV-1 infected dcOrgs rescued the ratios of extracellular Aβ42/40 to similar ratios as detected from uninfected dcOrgs. The results indicated that HSV-1 infection in dcOrgs led to a lower proportion of Aβ42 peptides that was secreted into the media.

### Expression of Iba1+ microglia and GFAP+ astrocytes were highly positively correlated with HSV-1

We next asked about the correlations in expression of HSV-1 with cell type specific marker expression using our flow cytometry approach. We observed positive correlations in the expression of HSV-1 and NeuN+ neurons (*r*=0.26 and 0.24; Figure S6), as well as positive correlations in the expression of HSV-1 and EOMES+ intermediate progenitor cells (*r*=0.3 and 0.44; Figure S6). Correlation analyses of glia cell-type marker intensities with HSV-1 intensities pointed to Iba1+ microglia with the highest correlations (*r*=0.66 and 0.39; Figures S7-8). There were no correlations observed in the expression of HSV-1 and P2RY12+ microglia (*r*=-0.05 and 0.09; Figure S7). We also observed positive correlations in the expression of GFAP+ astrocytes with HSV-1 (*r*=0.32 and 0.49; Figure S7), and GLAST+ astrocytes with HSV-1 (*r*=0.49 and 0.19; Figure S7).

These results showed that increased intracellular Aβ42 and pTau-212 expression in HSV-1 infected dcOrgs was likely to be mediated by interactions between neuronal cells with glia cells such as microglia and astrocytes, thus recapitulating key features of a neuroinflammation model for AD.

### Cell type enrichment analyses on RNA-seq data indicate an enrichment of astrocytes and depletion of excitatory neurons in HSV-1 infected dcOrgs

Glia-neuronal morphological and functional changes that are important in AD, include astrogliosis (marked by increased proportions of GFAP+ reactive astrocytes) and neuronal loss^61,62^. We sought to evaluate differences in the proportions of cell types by performing cell type enrichment analyses from the dcOrg RNA-seq datasets using the Orgo-Seq method that we had reported previously^34^. Briefly, the Orgo-Seq method uses cell type specific transcripts identified from published single-cell RNA sequence (scRNA-seq) datasets to identify cell types that might be enriched or depleted from RNA-seq data. We utilized a scRNA-seq dataset from human post-mortem brain samples as a reference dataset^63^ for Orgo-Seq (Figures 4A-F) and found that astrocytes were computed to be enriched in *Inf-vs-Ctrl*, *Inf-vs-ACV* and *Inf-vs-UV dcOrgs* (Figure 4A), which indicates that HSV-1 infection may result in increased proportions of astrocytes. On the other hand, excitatory neurons were computed to be depleted in *Inf-vs-Ctrl*, *Inf-vs-ACV* and *Inf-vs-UV dcOrgs* (Figure 4B), and similar reports of neuronal loss after HSV-1 infection had been previously observed^64–66^.

**Figure 4:**
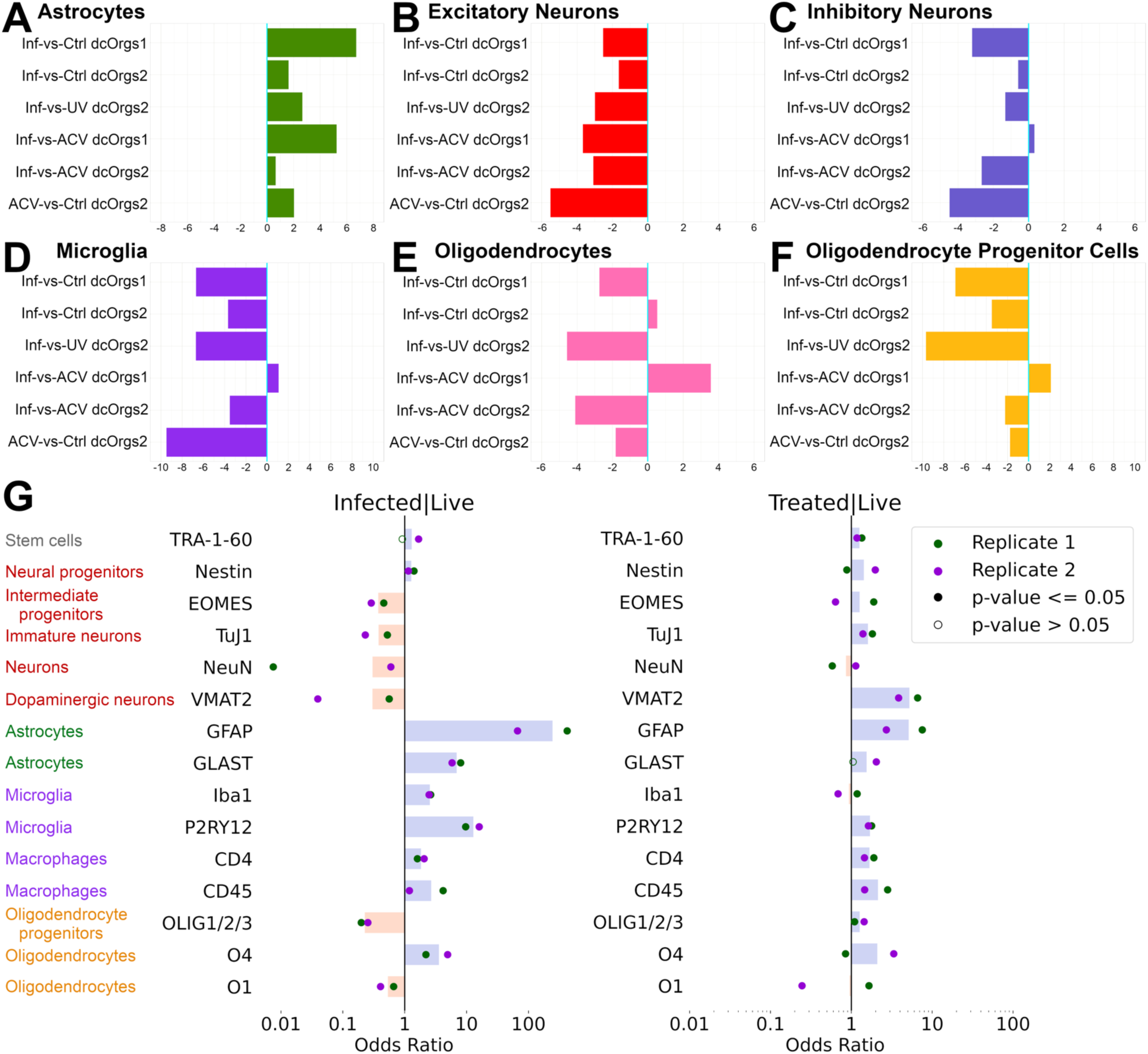
Cell-type specific analyses of HSV-1 infection and rescue by ACV. (A-F) *CellScores* calculated using Orgo-Seq method and *CellScores*>1 indicate that there is an enrichment, whereas *CellScores*<1 indicate that there is a depletion, of (A) astrocytes, (B) excitatory neurons, (C) inhibitory neurons, (D) microglia, (E) oligodendrocytes and (F) oligodendrocyte progenitor cells. (G) ORs calculated for HSV-1 infected live dcOrgs versus with uninfected live dcOrgs, or ACV-treated live dcOrgs versus uninfected live dcOrgs, using 14 cell-type markers. 2 sets of replicate experiments were performed for each marker (depicted by green and purple points). Shaded points represent significant odds ratios with *P*≤0.05 and open points represent non-significant odds ratios with *P*>0.05. The yellow bars represent average ORs<1 across both replicates and the blue bars represent average ORs>1 across both replicates.

### Flow cytometry results validated neuronal loss and astrogliosis observed from transcriptomics data, and provided insights into cellular biomarkers

To quantitatively evaluate cell type proportion changes in dcOrgs, we performed flow cytometry with 15 cell type specific antibodies. Using the cell counts identified from the flow cytometry results, we found that there were decreased proportions of live neurons and neural progenitor cells in HSV-1 infected dcOrgs compared to uninfected dcOrgs (EOMES+ intermediate progenitors, TuJ1+ immature neurons, NeuN+ mature neurons and VMAT2+ dopaminergic neurons), which indicated that there was neuronal death in dcOrgs due to HSV-1 infection (Infected live population in Figure 4G). Live glia and glia progenitor cells were mostly increased in proportions, such as GFAP+ or GLAST+ astrocytes, Iba1+ or P2RY12+ microglia, and O4+ oligodendrocytes. These results were consistent with reactive astrogliosis, activation of microglia and neuronal death previously observed in AD patients^67–69^.

ACV treatment of HSV-1 infected dcOrgs reduced the depletion in proportions of neurons and neural progenitor cells, and reduced the increased proportions of reactive glia cells (Treated live population in Figure 4G). CD4+ or CD45+ monocytes/macrophages did not show a huge increase in proportions among infected live cells and the proportions were similar after ACV treatment, which showed that the increased proportions of Iba1+ or P2RY12+ cells were driven by microglia and not monocytes nor macrophages.

### HSV-1 infection induced differentially expressed genes in dcOrgs that were enriched for AD-associated genes but not in hiPSCs or SH-SY5Y-differentiated neurons

We obtained lists of genes that were associated with 21 common neurodegenerative, neuropsychiatric or autoimmune diseases, as well as human traits and related diseases through genome-wide association studies (GWAS) from the GWAS Catalog^70^. To evaluate if the DEGs in infected versus uninfected dcOrgs were enriched for genes associated with any of the 21 common diseases or traits, we conducted gene set enrichment analyses (GSEA)^71,72^ with the gene lists in Table S6.

We observed that both sets of DEGs in *Inf-vs-Ctrl dcOrgs1* and *Inf-vs-Ctrl dcOrgs2* were enriched for AD-associated genes (*P*=0.039 and *P*=5.1×10^-3^ respectively; Figure 5A, Table S7). No enrichment was observed for the DEGs in *Inf-vs-Ctrl dcOrgs1* and *Inf-vs-Ctrl dcOrgs2* with genes associated with the other common diseases, suggesting that HSV-1 infection in dcOrgs is specifically perturbing the expression of AD-associated genes and gene networks. Similarly, the DEGs in HSV-1 infected dcOrgs versus dcOrgs with UV-inactivated HSV-1 (*Inf-vs-UV dcOrgs2*) were enriched for AD-associated genes (*P*=0.015), providing further support that the transcriptomic profiles of dcOrgs with UV-inactivated HSV-1 were similar to uninfected dcOrgs.

**Figure 5:**
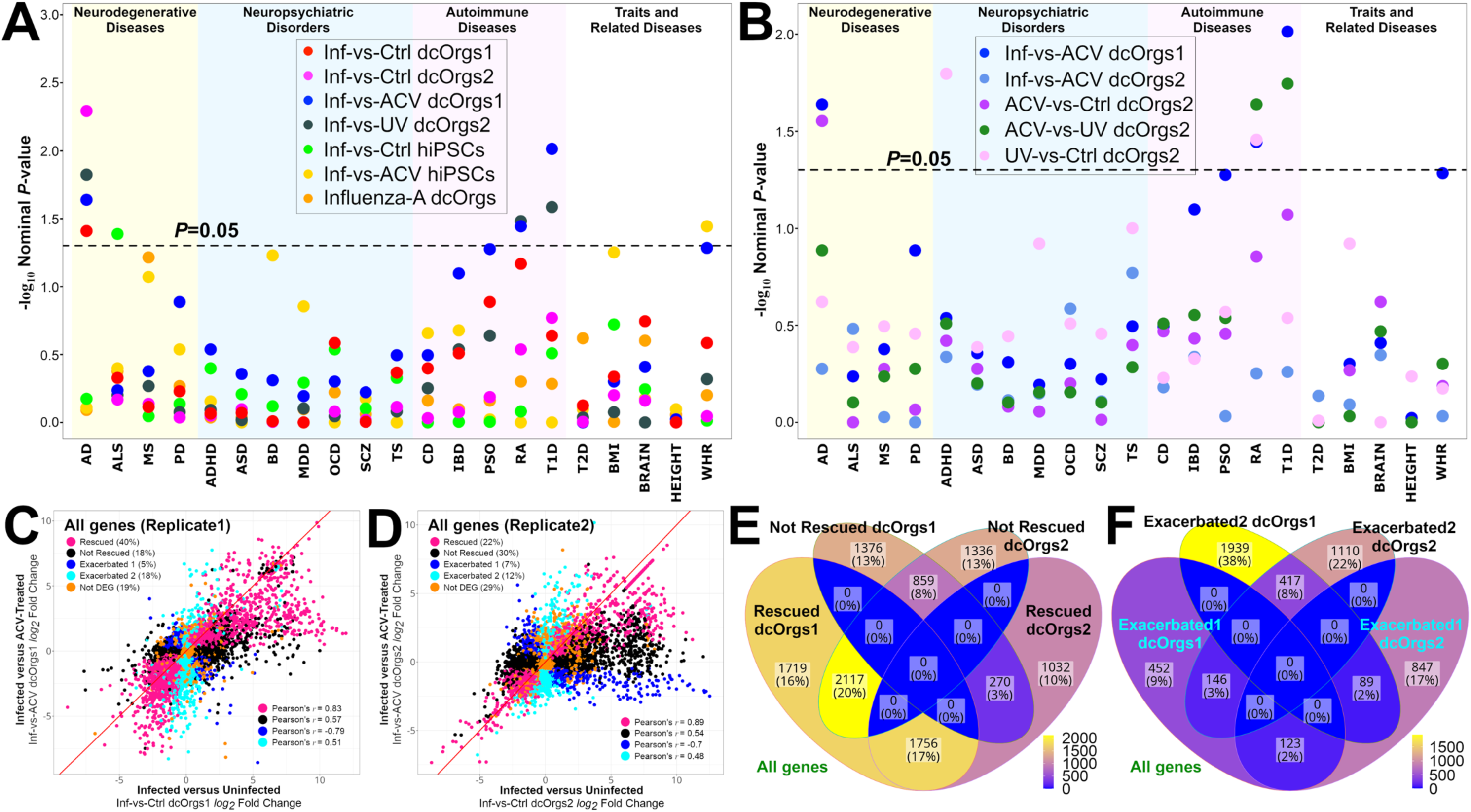
Analyses of DEGs from HSV-1-infected and ACV-treated samples. (A) -log_10_(Nominal *P*-value) from GSEA (*y*-axis) of the differentially expressed genes from our datasets with GWAS-associated genes in common diseases and traits (*x*-axis). (B) -log_10_(Nominal *P*-value) from GSEA (*y*-axis) of the differentially expressed genes from our datasets with GWAS-associated genes in common diseases and traits (*x*-axis). (C) Scatter plot for our discovery dataset showing the log_2_ fold change in infected versus ACV-treated dcOrg samples, *Inf-vs-ACV dcOrgs1* (*y*-axis) and infected versus uninfected dcOrg samples, *Inf-vs-Ctrl dcOrgs1* (*x*-axis), with the Pearson’s *r* correlations of the log_2_ fold changes of transcripts in each category shown on the bottom right. (D) Scatter plot for our replication dataset showing the log_2_ fold change in infected versus ACV-treated dcOrg samples, *Inf-vs-ACV dcOrgs2* (*y*-axis) and infected versus uninfected dcOrg samples, *Inf-vs-Ctrl dcOrgs2* (*x*-axis), with the Pearson’s correlations (*r*) of the log_2_ fold changes of transcripts in each category shown on the bottom right. (E) Venn diagram showing the numbers and percentages of genes that were rescued or not rescued in both our discovery and replication datasets (dcOrgs1 and dcOrgs2). (F) Venn diagram showing the numbers and percentages of genes that were in the exacerbated 1 or exacerbated 2 categories in both our discovery and replication datasets (dcOrgs1 and dcOrgs2).

We evaluated whether the DEGs in infected versus uninfected hiPSCs might be enriched for AD-associated genes, but we did not observe any enrichment (*P*=0.67; Figure 5A). We conducted GSEA using a published list of DEGs identified from HSV-1 infected versus uninfected SH-SY5Y-differentiated neurons^73^, but we did not observe any enrichment in AD-associated genes for the neuronal DEGs (*P*=0.38; Figure S9A). The results indicate that several of these transcriptomic perturbations in AD-associated genes are likely to be driven by type 1 interferon response produced by cell types other than stem cells or neurons within the dcOrgs^55,56,74–77^, such as glia cells.

### DEGs from IAV-infected dcOrgs were not enriched for AD-associated genes

We wondered whether inflammation induced by an RNA virus such as IAV, might similarly perturb AD-associated genes and therefore, we performed GSEA on DEGs identified from IAV-infected versus uninfected dcOrg samples. However, we did not observe an enrichment for AD-associated genes (*P*=0.79; Figure 5A, Figure S9B). This showed that HSV-1 infection, but not IAV infection, perturbed the expression of many AD-associated transcripts in the dcOrgs, potentially through the cytosolic DNA-sensing cGAS/STING/IRF3 pathways^78,79^ or other immune response pathways.

### ACV treatment in HSV-1 infected dcOrg samples rescued the expression in AD-associated genes

Next, we evaluated if ACV treatment of HSV-1 infected dcOrgs could rescue the transcriptomic perturbations in AD-associated genes that we had observed. GSEA analyses of our discovery dataset (*Inf-vs-ACV dcOrgs1*) showed an enrichment for AD-associated transcripts (*P*=0.023; Figure 5B), indicating that ACV treatment could rescue HSV-1 induced transcriptomic perturbations in AD-associated genes. However, our replication dataset (*Inf-vs-ACV dcOrgs2*) did not show an enrichment for AD-associated transcripts, indicating that ACV treatment in our replication experiment did not rescue HSV-1 induced transcriptomic perturbations in AD-associated genes (*P*=0.53; Figure 5B). As there were 14-20 times more HSV-1 viral transcripts detected in our replication dataset compared to our discovery dataset (Figure 1D), we hypothesized that there might be different extents of inhibition of HSV DNA synthesis in the two experiments.

Using a previously reported annotation of viral transcripts^80^, we observed that there was a high correlation in the *P*-value ranked differential expression for the leaky late viral transcripts (ψ_1_) across both datasets (Spearman’s ρ=0.63, *P*=1.6×10^-3^), but not for the true late viral transcripts (ψ_2_) across both datasets (Spearman’s ρ=0.2, *P*=0.47; Figure S10, Table S8). These results were consistent with prior reports that true late (ψ_2_) viral gene expression is perturbed by ACV treatment, but not immediate early, early or leaky late (ψ_1_) viral gene expression^80,81^. The results also indicate that our observed enrichment of AD-associated genes is due to expression of ψ_2_ viral genes in a dosage-dependent manner, indicating that the expression of one or more ψ_2_ genes led to differential expression in many of the AD-associated genes.

We performed GSEA using the DEGs from *ACV-vs-Ctrl dcOrgs2* and observed an enrichment for AD-associated genes (*P*=0.028; Figure 5B), which showed that the transcriptomic profiles of ACV-treated dcOrgs2 were more similar to infected dcOrgs and thus, the results from *ACV-vs-Ctrl dcOrgs2* were similar to *Inf-vs-Ctrl dcOrgs2* and showed a similar enrichment for AD-associated genes. These results indicated that ACV treatment can rescue transcriptomic perturbations in AD-associated DEGs due to HSV-1 infection by a mechanism that is dependent on inhibition of viral DNA synthesis.

Gene ontology analyses showed that biological processes such as regulation of apoptosis, autophagy and mitochondrion organization were enriched among DEGs in both sets of *Inf-vs-Ctrl dcOrgs* and *Inf-vs-ACV dcOrgs1* but not in *Inf-vs-ACV dcOrgs2* (Table S9). Pathways such as integrin signaling pathway, gonadotropin-releasing hormone receptor pathway and angiogenesis were enriched among DEGs in both sets of *Inf-vs-Ctrl dcOrgs* and *Inf-vs-ACV dcOrgs1* but angiogenesis was not an enriched pathway in *Inf-vs-ACV dcOrgs2* (Table S10).

### ACV treatment in HSV-1 infected dcOrgs led to enrichment of DEGs in genes associated with common autoimmune diseases

We observed that there were DEGs from some dcOrg samples that showed enrichment in genes associated with common autoimmune diseases such as Type 1 diabetes (T1D) and rheumatoid arthritis (RA). The DEGs from *Inf-vs-ACV dcOrgs1*, *Inf-vs-UV dcOrgs2* and *ACV-vs-UV dcOrgs2* showed enrichment in genes associated with both T1D and RA, whereas the DEGs from *UV-vs-Ctrl dcOrgs2* showed a similar enrichment in RA and attention-deficit hyperactivity disorder (ADHD) associated genes (Figures 5A-B). The enrichment of DEGs in T1D, RA and ADHD associated genes were not observed in *Inf-vs-Ctrl dcOrgs1* and *Inf-vs-Ctrl dcOrgs2*. These observations pointed to unexpected transcriptomic dysregulation of genes associated with autoimmune diseases by HSV-1 or ACV treatment, or capsid proteins found in UV-inactivated HSV-1, or a combination of these factors.

### ACV treatment exacerbated the expression for 19-23% of human transcripts that were expressed in dcOrgs

To explore the contribution of unexpected transcriptomic perturbations due to ACV treatment, we defined two categories of unexpectedly perturbed transcripts. First, we defined “Exacerbated 1” group as genes whose dysregulation were exacerbated by ACV treatment. For instance, if HSV-1 infection in dcOrgs down-regulated the human gene expression compared to uninfected dcOrgs, but ACV treatment in infected dcOrgs further down-regulated the gene expression compared to infected dcOrgs, or the reciprocal scenario with up-regulation, the gene would be classified into the Exacerbated 1 group. Next, we defined “Exacerbated 2” group as genes where HSV-1 infection in dcOrgs did not perturb the human gene expression but ACV treatment in infected dcOrgs significantly perturbed the expression of the gene.

Collectively, both groups of exacerbated genes comprised of 19% or 23% among all genes that were expressed in dcOrgs (Figures 5C-D). We observed that exacerbated genes comprised of similarly high percentages of 22% or 23% among AD-associated GWAS genes (Table S11). On the other hand, the expression for most of the genes (22% or 40%) were rescued by ACV treatment, and similarly 26% or 41% of AD-associated genes were rescued by ACV treatment (Table S11). These results reaffirmed that ACV treatment can rescue most AD-associated transcriptomic perturbations due to HSV-1 infection, despite resulting in high percentages of exacerbated gene expression.

### ACV treatment preferentially rescued AD-associated gene expression in dcOrg samples

We further explored the overlaps between transcripts that were rescued or not rescued across the dcOrg replicates. The highest overlap in transcripts were rescued in dcOrgs1 but were not rescued in dcOrgs2 (Figure 5E), which further supported our observations that ACV treatment was more effective in the discovery experiment (*Inf-vs-ACV dcOrgs1*) than the replication experiment (*Inf-vs-ACV dcOrgs2*). As expected, the second highest overlap in transcripts were rescued in both dcOrgs1 and dcOrgs2, demonstrating that despite the dosage-dependent differences of ACV treatment in both sets of experiments, there were several transcripts in common that were rescued across both datasets.

The highest overlap in AD-associated transcripts were driven by transcripts that were rescued in both dcOrgs1 and dcOrgs2 (Figure S11). There was an enrichment in the ratios of AD-associated transcripts that were rescued by ACV treatment, compared to the overall ratios of all transcripts that were rescued by ACV treatment (*OR*=1.33, *P*=0.062), suggesting that ACV treatment may have a preferential rescue for AD-associated genes in HSV-1 infected dcOrgs.

Similarly, there were modest enrichment of RA-associated or T1D-associated transcripts that were rescued in both dcOrgs1 and dcOrgs2, compared to the overall set of all transcripts (*OR*=1.33 and 1.47, *P*=0.068 and 0.074 respectively, Figure S11). There were 7 other diseases or traits with modest enrichments of transcripts that were rescued in both dcOrgs1 and dcOrgs2 (Figure S11), such as Parkinson’s Disease (PD; *OR*=1.55, *P*=0.032), ADHD (*OR*=1.24, *P*=0.071), obsessive compulsive disorder (OCD; *OR*=1.96, *P*=0.0043), schizophrenia (SCZ; *OR*=1.21, *P*=0.069), inflammatory bowel disease (IBD; *OR*=1.29, *P*=0.088), height (*OR*=1.14, *P*=0.03) and waist-hip-ratio (WHR; *OR*=1.34, *P*=7.3×10^-4^). These results indicate that ACV treatment may preferentially rescue the expression of AD-associated genes and genes associated with several other common diseases or traits.

On the other hand, the 4-way Venn diagrams depicting both groups of exacerbated genes did not show high overlaps in any subsets that were common between both dcOrg replicates (Figure 5F, Figure S12). This observation indicated that different sets of genes were transcriptionally exacerbated by each ACV treatment, unlike the rescued genes.

### HSV-1 and IAV infections in dcOrgs perturbed the expression of transcripts in several innate immune pathways

A strong transcriptional activation of the innate immune response had also been observed in our RNA-seq datasets from HSV-1 infection in dcOrgs and also reported previously from HSV-1 infection in 3D cerebral organoids^41^. This innate immune response is most likely produced by microglia or astrocytes within the cerebral organoids, although a small fraction (∼3%) of neurons had been reported to produce type I interferon response^77^. Neurons had also been reported to be activated directly by microbes to release neuropeptides and modulate the innate immune response^82,83^. As such, the interferon response observed from our data is likely to be primarily driven by glia cells (microglia, astrocytes) and possibly to a lesser extent, by neurons that differentiated innately within these dcOrgs.

The expression of transcripts in several human immune pathways (cGAS/STING, IFN, IRF3, JAK/STAT, RIG-I and TLR) were significantly differentially perturbed in both sets of HSV-1 infected versus uninfected dcOrgs (Figure 6A), as well as in hiPSCs (Figure S13). ACV treatment rescued the expression for some of these transcripts (Figure 6B). As expected, infected dcOrgs versus dcOrgs with UV-inactivated HSV-1 showed similar differential expression as infected versus uninfected dcOrg samples (Figure 6C). IAV-infected versus uninfected dcOrgs did not result in many significant DEGs in these human immune pathways, except in the RIG-I and TLR pathways (Figure 6D). These results are consistent with a recent report that the cGAS/STING pathway is a major driver of aging-related inflammation in neurodegenerative diseases such as AD^84^.

**Figure 6:**
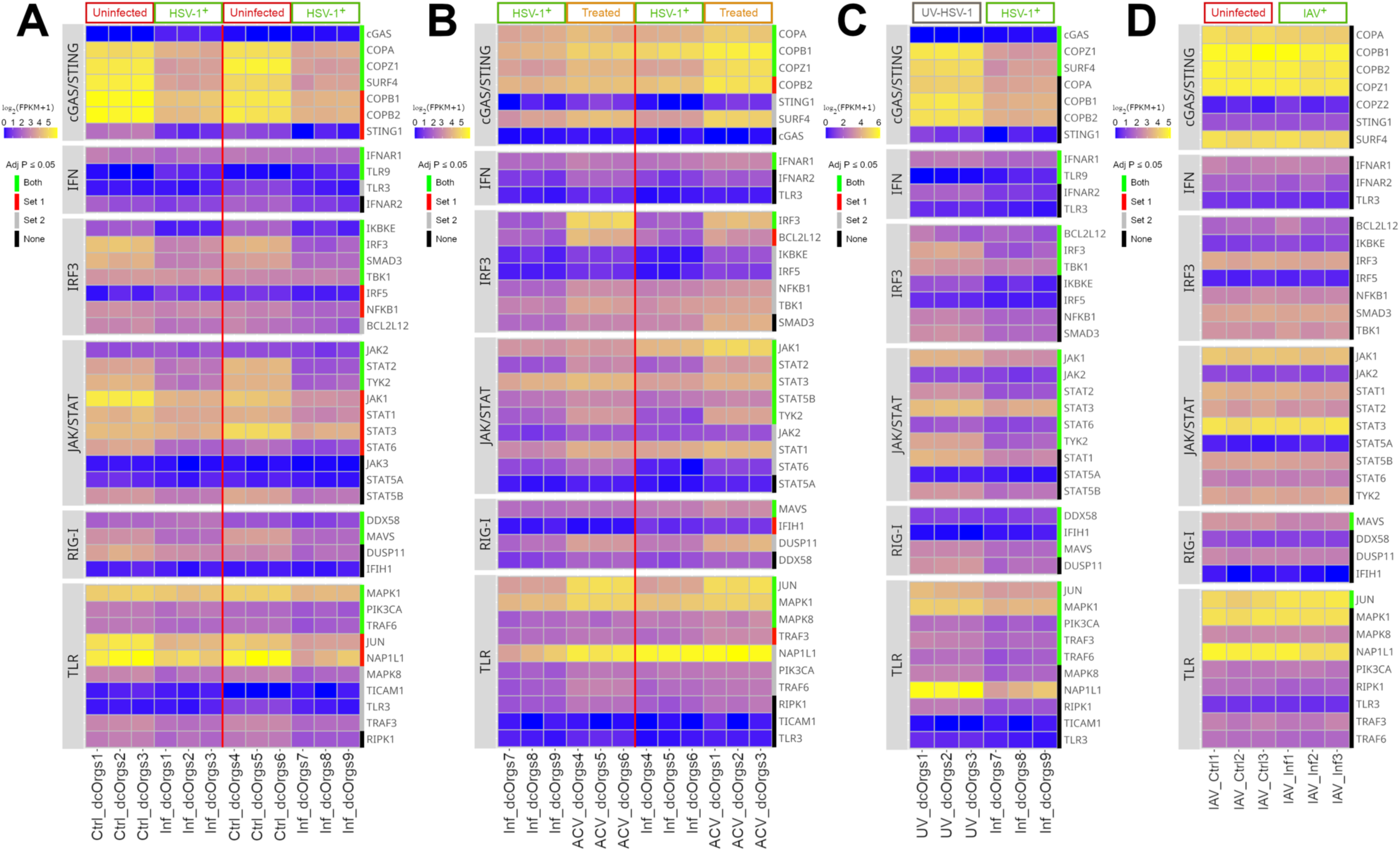
Heatmaps depicting the expressions of immune transcripts. (A) Heatmap depicting human transcript expression in HSV-1 infected or uninfected dcOrg samples. (B) Heatmap depicting human transcript expression from HSV-1 infected or ACV-treated dcOrg samples. (C) Heatmap depicting human transcript expression from dcOrg samples with UV-inactivated HSV-1 or HSV-1 infected dcOrg samples. (D) Heatmap depicting human transcript expression from IAV infected or uninfected dcOrg samples. (A-D) Green bars on the right of the heatmaps annotate transcripts that were significantly differentially expressed in both datasets (adjusted *P*≤0.05). Red bars on the right of the heatmaps annotate transcripts that were significantly differentially expressed in dcOrgs1 (adjusted *P*≤0.05) but not in dcOrgs2 (adjusted *P*>0.05). Gray bars on the right of the heatmaps annotate transcripts that were significantly differentially expressed in dcOrgs2 but not in dcOrgs1. Black bars on the right of the heatmaps annotate transcripts that were not significantly differentially expressed in both datasets (adjusted *P*>0.05). Red vertical lines in the heatmaps separate both sets of discovery and replication datasets.

### Analyses on human post-mortem brain RNA-seq revealed that 25-31% of patients with late-onset AD have transcriptomic signatures similar to HSV-1 infected dcOrgs

A previous study used RNA-seq data generated using human post-mortem brain samples from patients with late-onset AD (LOAD) and performed molecular subtyping of the LOAD patients into 5 subtypes (A, B1, B2, C1 and C2)^85^. Globally, there were weaker fold change differences observed in RNA-seq data from human post-mortem brains versus dcOrgs (Figure S14A). For better visualization, we rescaled the fold change differences observed in human post-mortem brains to illustrate that there were some pathways with high degrees of similarities in fold changes between the human post-mortem brains and dcOrgs, such as Amyloid, Blalock, Immune, Synapse/Myelin and Tau (Figure S14B).

We conducted a modified gene set enrichment analysis test using the DEGs from the dcOrgs with the most informative DEGs from the post-mortem brains and identified a significant enrichment of the DEGs in dcOrgs with subtype A (Figure S14C, Table S12). *Inf-vs-Ctrl dcOrgs1*, *Inf-vs-Ctrl dcOrgs2*, *Inf-vs-ACV dcOrgs1* and *ACV-vs-Ctrl dcOrgs2* had significant positive associations with subtype A (*FWER*=1×10^-8^, 9.6×10^-9^, 6.4×10^-10^ and 4×10^-7^ respectively). Subtype A comprises of 25% and 31% of the LOAD patients from two cohorts^85^, suggesting that a substantial fraction of LOAD patients demonstrate molecular signatures that resemble transcriptomic signatures from our HSV-1-associated dcOrg neuroinflammation system for AD.

## DISCUSSION

Neuroinflammation had been proposed as a central mechanism in AD^2,86^. We tested the hypothesis that HSV-1 infection of human dcOrgs could result in *in-vitro* cellular and molecular changes associated with common neurodegenerative, neuropsychiatric or autoimmune diseases. We found that HSV-1 infected dcOrgs led to AD-associated pathologies, such as decreased ratios of secreted extracellular Aβ42/40, increased intracellular Aβ42 and pTau-212 expression, neuronal death, astrogliosis and activation of microglia, which are hallmarks of AD^62,87^, as well as transcriptomic perturbations that were enriched for AD-associated genes. Immediate treatment with ACV on HSV-1 infected dcOrgs rescued these cellular and molecular changes.

However, our study also found that ACV treatment may potentially result in unexpected transcriptomic perturbations of genes associated with autoimmune diseases such as RA and T1D. These unexpected transcriptomic perturbations may not lead to cellular or physiological effects.

Given the ongoing clinical trials for VCV as an AD therapeutics^22,23^, our findings can aid in understanding the safety and effect of anti-herpetic drugs for AD.

In addition, recent evidence indicates that the innate immune pathway cGAS/STING may be a major contributor in neurodegenerative diseases^84^. Our study similarly pointed to neuroinflammation by the human innate immune pathways due to DNA viruses such as HSV-1, which led to AD-associated cellular and molecular pathologies. Further research into the role of the innate immune pathways in AD may highlight mechanisms and therapeutic targets for reversing neuroinflammation due to neurotropic viruses or other pathogens.

We also found that post-mortem brain samples from 25% or 31% of LOAD patients have transcriptomic signatures that were similar to HSV-1 infected dcOrgs, suggesting that a major fraction of AD patients may benefit from anti-HSV-1 or more broadly, anti-herpetic treatment. Our study can also inform the selection of AD patient subtypes who may benefit from anti-viral therapeutics.

Our high-throughput, quantitative, *in-vitro* screening platform using Aβ42/40/38 ELISA assays, flow cytometry and transcriptomics with human-derived dcOrgs provides a high-throughput system that can be used for biomarker discovery, and to identify potential therapeutic targets and compounds that can rescue the cellular and molecular changes associated with AD due to neuroinflammation.

## EXPERIMENTAL PROCEDURES

### Standard Protocol Approval

Research performed on samples and data of human and viral origin was conducted according to protocols approved by the Institutional Review Board (IRB) and Institutional Biosafety Committee (IBC) of UMass Chan Medical School. HSV-1 and IAV are Biosafety Level (BSL) 2 pathogens.

### Source of hiPSCs and maintenance of hiPSCs

The control donor hiPSC line used in our study (PGP1 iPSC) was a kind gift from Professor George Church’s lab, and we had previously characterized and sequenced the line^34,88^. We performed flow cytometry (MACSquant VYB) to confirm that >90% of the PGP1 hiPSCs are positive for TRA-1-60 (Novus Biologicals NB100-730F488). hiPSCs were plated using StemFlex media (Thermo Scientific A3349401) with 10μM Y-27632 (Abcam ab120129) on 6-well plates coated with 0.5mg of Matrigel Basement Membrane Matrix (Corning 354234) per plate, and subsequently dissociated using accutase (BioLegend 423201).

### HSV-1 virus and ACV

The HSV-1 K26GFP KOS strain^89,90^ used in our study encodes a GFP-fused VP26 gene and was kindly provided by Prashant Desai^91^. We prepared 200mM stock solutions of ACV (Sigma PHR1254) by dissolving in UltraPure DNase/RNase-Free Distilled Water (Invitrogen 10977015) with equal molar concentrations of sodium hydroxide. Aliquots were made and kept frozen in - 80°C.

### Influenza A virus

The IAV virus (strain (A/Puerto Rico/8/1934(H1N1))) used in our study^92^ was purchased from Charles River Laboratories. The virus was propagated in SPF eggs in the allantoic cavity. Clarified allantoic fluid was concentrated and resuspended in Hepes-Saline solution and layered on a sucrose gradient. The interface band was diluted, pelleted and resuspended in a minimal volume of Hepes-Saline solution. Antigen was tested for protein concentrate of 2mg of protein per mL using a Bio-Rad colorimetric protein assay. The final HA titer per 0.05mL was 131,072 and the EID50 titer per mL was 10^9.8^. A FITC-conjugated anti-Influenza A NP monoclonal antibody (ThermoFisher MA1-7322) at 1:20 dilution was used to detect viral proteins using flow cytometry.

### Cerebral organoid differentiation

We adapted a previously described protocol to generate spontaneously differentiated dcOrgs from PGP1 hiPSCs^34,93,94^. We re-suspended 900,000 cells in 15ml of StemFlex with 50μM Y-27632 and seeded the cells across a 96-well ultra-low attachment plate (Corning CLS7007) to form embryoid bodies. After 5 days, each embryoid body was transferred to 24-well ultra-low attachment plates (Corning CLS3473) with 500μL of neural induction media in each well. An additional 500μL of neural induction media was added to each well on Day 8. On Day 10, each organoid was embedded in 40μL of Matrigel (Corning 354234) on parafilm and incubated at 37°C for 15 minutes before they were scraped into the wells containing 2mL of differentiation media using a cell scraper.1-2mL of differentiation media with 10% penicillin streptomycin (ThermoFisher 15140122) per well was used to passage the organoids every 3-7 days, and the plates of organoids were placed on an orbital shaker at 90rpm.

### Cerebral organoid dissociation

After 2-4 months of differentiation, we dissociated 168-192 dcOrgs in each batch and dcOrgs were washed twice in ice-cold 1×DPBS (ThermoFisher 14190144) for 10 minutes at 4°C, followed by incubation in 500μL of 0.25% Trypsin-EDTA (ThermoFisher 25200056) for 15 minutes at 37°C and 300rpm with repeated manual pipetting to dissociate clumps of cells^94^. To remove cell debris, we further filtered the cells using a 30μm MACS SmartStrainer (Miltenyi Biotec 130-098-458). Cell counts and viabilities were measured using an automated cell counter (NanoEnTek EVE). Cell viabilities ranged from 80% to 96%, and we obtained 7.8×10^-6^ to 4.1×10^-7^ live cells in total. Subsequently, the dissociated cells were passaged in differentiation media for at least a month to allow the cells to recover. We recorded the total time allowed for differentiation and recovery of the cells prior to infection as the age of the dcOrgs.

### HSV-1 and IAV infection experiments

For infections in 3D cerebral organoids, we first washed 3-month intact cerebral organoids twice (one organoid in a single 1.5mL tube) in sterile ice-cold 1×DPBS, followed by a 10-minute spin-down. We used MOI of 2 and 10 for HSV-1 infections in 3D cerebral organoids, and by assuming that there were 15,000 cells on the surface of the organoids. Subsequently, after 23 hours or 47 hours, we took microscope images of the uninfected and infected organoids. We also quantified the percentage of infected cells in the 3D cerebral organoids using flow cytometry. We pooled 20 uninfected or infected 3D cerebral organoids into 1.5mL tubes, washed twice using 1×DPBS, dissociated the organoids using 0.25% Trypsin-EDTA and fixed the cells with 4% paraformaldehyde (PFA; Fisher Scientific 50-980-487) for flow cytometry.

For HSV-1 infections in 2D dcOrgs, we infected 3-month dcOrgs using MOI of 1, 2, 4 or 10, and found that MOI of 2 resulted in >50% of infected dcOrgs that were positive for GFP expression, quantified using flow cytometry. We used a higher virus titer (MOI of 4) for hiPSCs. We counted the number of cells needed for each experiment and seeded the cells at a density of 1×10^6^ cells/well of a 6-well tissue culture plate coated with 0.5mg of Matrigel, using 2mL/well of differentiation media with 10μM Y-27632. On the following day, cells from one of the wells were counted using an automated cell counter to calculate the amount of virus needed.

Inoculation media containing 1×DPBS with 0.5% FBS (ThermoFisher 10082147) was prepared, and aliquots of HSV-1 and ACV were kept on ice. Cells in each well were washed with 2mL of 1×DPBS. 1mL of inoculation media was added to each well of uninfected control cells, 1mL of inoculation media with HSV-1 was added to each well of infected cells, and 1mL of inoculation media with HSV-1 and 200μM ACV was added to each well of treated cells. The cells were placed back into an incubator at 37°C for an hour, and each well of cells were subsequently washed with 2mL of 1×DPBS. 2mL of differentiation media was added to each well of uninfected or infected cells, and 2mL of differentiation media with 200μM ACV was added to each well of treated cells. The cells were placed back into an incubator at 37°C for 23 hours. We inspected the cells under a fluorescence microscope (Logos Biosystems CELENA S Digital Imaging System) to visually verify the presence of GFP in the infected and treated cells. A similar protocol was used for IAV infections, except that MOI of 3 was used with a 47-hour post-inoculation incubation.

### RNA extraction and sequencing

Cells were fixed using 4% PFA and RNA was extracted from the fixed cells using the PureLink FFPE RNA Isolation Kit (ThermoFisher K156002), and treated with Ambion DNase I (ThermoFisher AM2222) to digest genomic DNA according to the manufacturers’ protocols. Total RNA was shipped overnight on dry ice to Psomagen for quantification (Table S1), followed by ribosomal RNA depletion and library preparation (Illumina TruSeq Stranded Total RNA Library Prep 20020596), followed by 151bp paired-end sequencing with a total of 40 million reads on a NovaSeq6000.

### RNA sequence alignment and data processing

Quality control was performed with the FastQC tool^95^, and adapter sequences and low quality base calls were trimmed from FASTQs with Trimmomatic v0.39^96^. Trimmed FASTQs were aligned to a concatenated GRCh37 human reference genome, human alphaherpesvirus 1 Kos strain reference genome (GenBank: JQ673480.1) and influenza A strain A/Puerto Rico/8/1934(H1N1) reference genome (NCBI BioProject PRJNA485481) using HISAT2 v2.2.1^97^. SAMtools v1.9 was used to extract reads aligning to the coding sequences of the virus^98^. Expression quantification and preprocessing was performed using StringTie2 v1.3.6^99^.

The transcript list was filtered to include only protein coding transcripts and duplicate genes in the StringTie2 output were removed, resulting in a total of 19,163 unique genes. We compared 2 approaches for alignment: firstly, we aligned all transcripts to a custom reference comprised of both human and HSV-1 reference sequences and secondly, we aligned all transcripts to human and HSV-1 reference sequences individually. We calculated Pearson’s *r* between the raw gene expression using both alignment approaches and found high correlations between both approaches (1-*r*^2^≤1×10^-4^). As a quality control step, we checked that there was no HSV-1 viral transcript expression in all control uninfected samples. Pearson’s *r* were calculated using the fragments per kilobase of exon per million mapped fragments (FPKM) values between sample pairs.

### Differential expression analyses

We performed differential expression analyses using the DESeq2 v1.38.2 package in R^100^. Genes were filtered using a cutoff of counts per million (CPM)≥1 across at least 2 samples. Raw counts were normalized using DESeq2 TMM normalization and models subsequently generated on a categorical design of infection or treatment status. The adjusted *P*-values from the differential expression analyses used in our study were Benjamini-Hochberg adjusted *P*-values calculated from the raw *P*-values. Volcano plots were generated in R^101^ with EnhancedVolcano v1.11.3^102^ and heatmaps were generated in R^101^ with ggplot2^103^.

### Ratios of human transcripts that were up-regulated versus down-regulated

The ratios were calculated as the numbers of human transcripts post-quality control that were up-regulated with positive log-fold changes divided by the numbers of human transcripts post-quality control that were down-regulated with negative log-fold changes. Fisher’s Exact Test *P*-values were calculated, compared against a null where there are equal numbers of human transcripts with positive and negative log-fold changes. To account for multiple hypotheses testing, Bonferroni correction for *P*-values that were 0.1 was applied to obtain the *FWER* values.

### Antibody labeling and flow cytometry experiments

Cells were washed with 1×DPBS and pelleted into 1.5mL microcentrifuge tubes. Each cell pellet was resuspended in 100μL of 1×DPBS with 1:500 Zombie Violet (BioLegend 423113) and incubated for 20 mins at room temperature in the dark. The cell pellets were then washed with 500μL of cell staining buffer (BioLegend 420201) and resuspended in either 100μL of 4% PFA or 250μL of Cytofix/Cytoperm fixation and permeabilization buffer (BD Biosciences 554714), and incubated at 4°C for 20 mins at 300rpm. For intracellular antibodies, the cells were then washed once with 500μL of permeabilization solution. For cell surface antibodies, the cells were then washed with 500μL of cell staining buffer, then resuspended in 105μL of cell staining buffer with 1:21 Human TruStain FcX (BioLegend 422301), followed by incubation at 4°C for 20 mins at 300rpm. Each cell pellet was then resuspended in 50μL of the respective buffer with 1:21 Human TruStain FcX and the antibody of interest, followed by incubation at 4°C for 60 mins at 300rpm. After antibody labeling, each cell pellet was washed twice with 500μL of Cell Staining Buffer, resuspended in 300μL of Cell Staining Buffer and transferred to a 5mL glass tube. We used a BD

Biosciences FACSCelesta Cell Analyzer at the UMass Chan FACS core facility for our flow cytometry experiments. Using the FACSDiva software, a “usable cell” gate was drawn on the FSC-A/SSC-A plot to minimize the collection of debris, and 20,000-30,000 cells within the useable cell population were analyzed.

### List of antibodies used

The antibodies used in our study were: Alexa Fluor 647-conjugated Aβ1-42 (Bioss Antibodies bs-0107R-BF647) at a 1:50 dilution, Alexa Fluor 647-conjugated Tau (Thr212) (Bioss Antibodies bs-5420R-BF647) at a 1:50 dilution, Alexa Fluor 647-conjugated Solanezumab (Novus Biologicals FAB9919R) at a 1:50 dilution, Alexa Fluor 647-conjugated TRA-1-60 (BioLegend 330605) at a 1:20 dilution, Alexa Fluor 647-conjugated Nestin (Novus Biologicals IC1259R-100UG) at a 1:50 dilution, Alexa Fluor 647-conjugated EOMES (Novus Biologicals IC6166R-100UG) at a 1:50 dilution, APC-conjugated TuJ1 (Biolegend 801219) at a 1:20 dilution, Alexa Fluor 647-conjugated NeuN (Novus Biologicals NBP1-92693AF647) at a 1:50 dilution, Alexa Fluor 647-conjugated VMAT2 (R&D Systems FAB8327R) at a 1:20 dilution, Alexa Fluor 647-conjugated GFAP (BioLegend 644706) at a 1:20 dilution, APC-conjugated GLAST (Miltenyi 130-123-555) at a 1:50 dilution, Alexa Fluor 647-conjugated Iba1 (Novus Biologicals 603102) at a 1:50 dilution, APC-conjugated P2RY12 (BioLegend 392113) at a 1:20 dilution, Alexa Fluor 647-conjugated CD4 (Biolegend 300520) at a 1:20 dilution, Alexa Fluor 647-conjugated Olig1/2/3 (Bio-techne FAB2230R-MTO) at a 1:50 dilution, Alexa Fluor 647-conjugated O4 (R&D Systems FAB1326R) at a 1:20 dilution and Alexa Fluor 647-conjugated O1 (R&D Systems FAB1327R) at a 1:20 dilution.

Isotype controls in our study used the following antibodies: Alexa Fluor 647-conjugated rabbit IgG (Bioss Antibodies bs-0295P-A647) at a 1:50 dilution, Alexa Fluor 647-conjugated mouse IgG1 kappa (Novus Biologicals IC002R) at a 1:50 dilution, Alexa Fluor 647-conjugated mouse IgG2A kappa (Biolegend 400234F2) at a 1:20 dilution and Alexa Fluor 647-conjugated mouse IgG2B kappa (Novus Biologicals IC0041RF3) at a 1:20 dilution.

### Machine learning based gating for flow cytometry data

Flow cytometry data was exported from FlowJo v10 as a .csv file with the per cell values for the Alexa Fluor 488-A, Alexa Fluor 647-A, Pacific Blue-A, FSC-A, FSC-H, SSC-A, and SSC-H channel intensities. FSC/SSC channels were *log*_10_-transformed, and infinite, maximum or zero values were dropped from further analyses. Next, viable cells were identified based on FSC-A and SSC-A minimum and maximum cutoffs. Modality was determined by calculating the number of local maxima on the kernel density of the data with a height of at least 15% of the global maximum. For unimodal data, the minimum and maximum cutoffs were set at the 5th and 95th percentiles respectively. For multimodal data, the data was fit to a Bayesian Gaussian mixture model with n_components equal to the number of peaks, in order to separate the data into distinct distributions. Bimodal data was then gated at the 5th and 95th percentile of the distribution with the higher intensities, while multimodal data was gated at the 5th and 95th percentile of the distribution with the average intensities. Gates were calculated based on the uninfected sample and applied to the infected and ACV-treated samples.

We next tested for singlets by fitting a linear regression model using FSC-A versus FSC-H intensities for each condition (uninfected, infected, or treated). We calculated the residuals and cells with residuals greater than the 99th percentile were considered to be doublets and were removed from further analyses.

Pacific Blue-A, Alexa Fluor 488-A, and Alexa Fluor 647-A gate cutoffs were calculated similarly to FSC/SSC gates. First, the modality was determined by calculating the number of local maxima on the kernel density of the data with a height of at least 15% of the global maximum, and distributions were separated using a Bayesian Gaussian mixture model with n_components equal to the number of peaks. Pacific Blue-A and Alexa Fluor 488-A gates were based on the infected sample distribution, while Alexa Fluor 647-A gates were based on the uninfected sample distribution, and the same gates were applied to the other samples. If the selected distribution was not bimodal, but the ACV-treated distribution was bimodal, then the gate was set based on the ACV-treated distribution and applied to the other samples. The cutoff for unimodal data was set at the 85th percentile, while the cutoff was set at the 1st percentile of the distribution with the higher intensities for bimodal data and the 85th percentile of the distribution with average intensities for multimodal data.

Bayesian Gaussian mixture models were implemented using the Scikit-learn python package, kernel density plots were calculated using the seaborn python package, and peaks were calculated using the SciPy python package.

### UMAP plots

UMAP was performed using the Python package umap-learn. For each flow experiment, the initial UMAP values for Alexa Fluor 488-A, Alexa Fluor 647-A, and Pacific Blue-A were normalized between 0 and 1 across all conditions (uninfected, infected and ACV-treated). The data for these three channels were then fitted to a UMAP model initialized with the number of components = 2 to re-calculate UMAP1 and UMAP2 values.

### Adjusted correlations in Alexa Fluor 488 and Alexa Fluor 647 intensities

We first performed linear regression correlation using the Alexa Fluor 488 and Alexa Fluor 647 intensities from the uninfected samples, similar to calculating the Pearson’s correlation *r*. This enabled us to identify the background correlation in the intensities. We obtained the *β* coefficient from the linear regression model and corrected the Alexa Fluor 647 intensities (*y_adjusted_* or adjusted *r*) for the infected and ACV-treated correlations as follows:

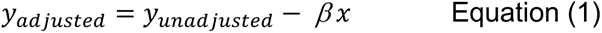

where *y_adjusted_* were the unadjusted Alexa Fluor 647 intensities in the infected and ACV-treated data and *x* were the Alexa Fluor 488 intensities in the infected and ACV-treated data.

To evaluate if the normalized Alexa Fluor 647 intensity distributions were lower in the uninfected versus infected cells within the same infected/ACV-treated sample, we calculated 1-tailed Wilcoxon ranked sum tests (the *P*-values in the boxen plots).

### Odds ratio calculations from flow cytometry data

Using the machine learning based gating, we grouped the cells into 4 quadrants: Q1 where the cells were negative for HSV-1 (GFP) and positive for the marker of interest (Alexa Fluor 647 or APC), Q2 where the cells were positive for both HSV-1 and the marker of interest, Q3 where the cells were positive for HSV-1 and negative for the marker of interest, and Q4 where the cells were negative for both HSV-1 and the marker of interest. We calculated ORs and Fisher’s Exact Test *P*-values in R^101^, where:

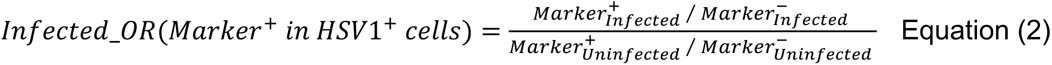

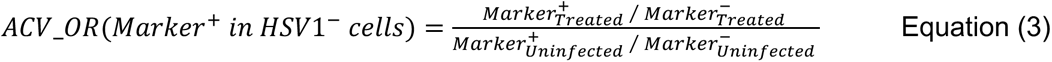

### ELISA assays and analyses

After inoculation, 1mL of fresh dcOrg differentiation media was dispensed into each well and cells were incubated for 48 hours. Conditioned media from each well were collected into 2 tubes with 500μL of media in each tube. Each set of conditions (uninfected dcOrg controls for HSV-1, HSV-1 infected dcOrgs, ACV-treated dcOrgs, uninfected dcOrg controls for IAV and IAV infected dcOrgs) had 4-5 experimental replicates. One set of conditioned media was heat inactivated at 65°C for an hour and the second set of conditioned media was UV-inactivated. We measured extracellular Aβ42/40/38 levels were measured using the V-PLEX Plus Aβ Peptide

Panel 1 (6E10) Kit (MSD K15200G-1), with controls diluted in the given buffer^27^. The Aβ peptide values were batched normalized to control in the ELISA kit. The concentrations of each Aβ peptide and ratios of Aβ peptides were visualized using ggplot^103^ in R^101^. 1-tailed Wilcoxon ranked sum tests were used to evaluate the samples of interest versus matched uninfected controls.

### Gene lists from GWAS catalog and gene set enrichment analyses

Gene lists associated with 21 common diseases were downloaded from the NHGRI-EBI GWAS catalog^70^ in April 2022. Gene set enrichment analyses were performed using GSEA v4.3.2^71,72^ to determine the enrichment of DEGs ranked by absolute *log2* fold change in decreasing order with the GWAS-associated gene lists. Analyses were performed using 10,000 permutations and a weighted enrichment statistic.

### Definitions of rescued, not rescued, exacerbated 1 and 2 transcripts

Rescued transcripts were defined as transcripts that were significantly differentially expressed (*P*≤0.05) in both the *Inf-vs-Ctrl dcOrgs* and *Inf-vs-ACV dcOrgs* datasets and *log_2_* fold changes in both datasets were in the same direction. For instance, a positive *log_2_* fold change in *Inf-vs-Ctrl dcOrgs* indicated that HSV-1 infection led to increased expression of the transcript. Similarly, a positive *log_2_* fold change in *Inf-vs-ACV dcOrgs* meant that ACV treatment of HSV-1 infected dcOrgs lowered the expression of the transcript, with respect to the expression in infected dcOrgs.

“Not rescued” transcripts were defined as transcripts that were significantly differentially expressed (*P*≤0.05) in *Inf-vs-Ctrl dcOrgs* but were not significantly differentially expressed (*P*>0.05) in Inf-vs-ACV dcOrgs.

“Exacerbated 1” transcripts were defined as transcripts that were significantly differentially expressed (*P*≤0.05) in both the *Inf-vs-Ctrl dcOrgs* and *Inf-vs-ACV dcOrgs* datasets but *log_2_* fold changes in both datasets were in the opposite direction. For instance, a positive *log_2_* fold change in *Inf-vs-Ctrl dcOrgs* indicated that HSV-1 infection led to increased expression of the transcript. However, a negative *log_2_* fold change in *Inf-vs-ACV dcOrgs* meant that ACV treatment of HSV-1 infected dcOrgs further increased the expression of the transcript, with respect to the expression in infected dcOrgs.

“Exacerbated 2” transcripts were defined as transcripts that were not significantly perturbed in *Inf-vs-Ctrl dcOrgs* (*P*>0.05) but were significantly differentially expressed in *Inf-vs-ACV dcOrgs* (*P*≤0.05). This indicates that ACV treatment on HSV-1 infected dcOrgs significantly perturbed the expression of these transcripts while the expression of these transcripts were not perturbed by HSV-1 infection in dcOrgs, with respect to the expression in uninfected dcOrgs.

### Scatter plots to represent rescued, not rescued, exacerbated 1 and 2 transcripts

Scatter plots showed the *log_2_* fold changes in the *Inf-vs-Ctrl dcOrgs* versus the *log_2_* fold changes in *Inf-vs-ACV dcOrgs* for all transcripts that were in common between both datasets. The percentages of transcripts in each category (rescued, not rescued, exacerbated 1, exacerbated 2 or not DEG) were calculated as the number of transcripts in each category divided by the total number of transcripts and multiplied by 100%. Pearson’s *r* were calculated between the *log_2_* fold changes in *Inf-vs-Ctrl dcOrgs* and *Inf-vs-ACV dcOrgs* for the transcripts in each category.

### Enrichment of disease-associated rescued or exacerbated transcripts

Fisher’s Exact Test was used to calculate the enrichment of the number of disease-associated transcripts in a group (such as the transcripts that were rescued in both dcOrgs1 and dcOrgs2) over the number of disease-associated transcripts that were not in the group, compared to (the total number of transcripts in the group minus the number of disease-associated transcripts in the group) over (the total number of transcripts that were not in the group minus the number of disease-associated transcripts that were not in the group).

### Enrichment analyses of AD subtype and dcOrg DEGs

We used the results from an earlier study that identified subtypes using LOAD post-mortem brain samples in the Mount Sinai/JJ Peters VA Medical Center Brain Bank (MSBB-AD) and Religious Orders Study-Memory and Aging Project (ROSMAP) cohorts^85^. We calculated the cumulative overlap fraction of DEGs for each of the 5 subtypes, which we defined as 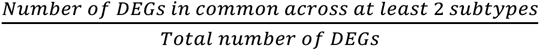 cumulative overlap fractions for each subtype to identify that the points of inflection across all subtypes converge at ∼200 DEGs, indicating that the top 200 DEGs across the 5 subtypes were likely to be most informative for subtype distinction.

Gene set enrichment analyses of the dcOrg DEGs ranked by decreasing order of absolute *log_2_* fold changes and the ranked DEGs from each subtype were calculated using connectivity scores^20^. Enrichment scores were permuted 10,000 times to determine the significance of the observed scores.

## ACKNOWLEDGEMENTS

We thank all members of the Accelerating Medicines Partnership Program for Alzheimer’s Disease (AMP-AD) group and all members of the AMP-AD working groups (Experimental Validation, Multi-scale Analysis) for their constructive feedback on the research. We also thank all members in the Departments of Genomics and Computational Biology (GCB); Neurology; and Molecular Cell and Cancer Biology (MCCB), especially the late Dr. Michael Green, for their constructive scientific feedback. This study was supported by the National Institutes of Health (NIH) AMP-AD grant U01AG061835 (PI: B.R.; Sub-contract PIs: G.M.C., E.T.L., Y.C.), NIH P01 grant AI09681 (PI: D.M.K), and UMass Chan Medical School startup funds (PIs: E.T.L., Y.C.).

## AUTHOR CONTRIBUTIONS

E.T.L. and Y.C. conceived the project. P.D., N.J.B., Y.C. and E.T.L. designed and performed the computational analyses and methods development. M.N.O., L.F.M., J.S., Y.C. and E.T.L. designed and performed the cerebral organoid differentiation and dissociation, viral infections for RNA extraction and quality control. H.S.O., Y.C. and D.M.K. designed and performed the initial HSV-1 experiments. H.S.O. and D.M.K. provided stocks of viruses and acyclovir. M.N.O., L.F.M, J.S., A.R.O., S.M.C., K.T., K.A., M.G., Y.C. and E.T.L. designed and performed the flow cytometry experiments and analyses. L.F.M., A.J.A., T.L.Y-P. and E.T.L. designed and performed the ELISA experiments. M.F.C., S-P.H.R., Q.W., F.H., M.H.O., D.G., M.H., G.M.C., B.R. and D.M.K. provided datasets and guided the analyses related to virology, immunology and technologies. G.M.C., B.R., D.M.K., Y.C. and E.T.L. wrote the grants to provide funding for the research. E.T.L. wrote the initial manuscript with edits from all authors.

